# LANTSA: Landmark-based transferable subspace analysis for single-cell and spatial transcriptomics

**DOI:** 10.1101/2022.03.13.484116

**Authors:** Chuanchao Zhang, Lequn Wang, Xinxing Li, Wei-Feng Guo, Qianqian Shi, Luonan Chen

## Abstract

Single-cell RNA sequencing (scRNA-seq) and spatial transcriptomics (ST) technologies provide new insights to understand tissue organization and biological function. Accurately capturing the relationships of samples (e.g., sequenced cells, spatial locations) will result in reliable and consistent outcomes in downstream analyses. However, this undertaking remains a challenge for large-volume or cross-platform datasets due to transcriptional heterogeneity and high computational demands. Here, we introduce landmark-based transferable subspace analysis (LANTSA) to solve such challenges for scRNA-seq and ST datasets. Specifically, LANTSA constructs a representation graph of samples for clustering and visualization based on a novel subspace model, which can learn a more accurate representation and is theoretically proven to be linearly proportional to data size in terms of the time consumption. Furthermore, LANTSA uses a dimensionality reduction technique as an integrative method to extract the discriminants underlying the representation structure, which enables label transfer from one (learning) dataset (i.e., scRNA-seq profiles) to the other (prediction) datasets (e.g., scRNA-seq or ST profiles), thus solving the massive-volume or cross-platform problem. We demonstrated the superiority of LANTSA to identify accurate data structures via clustering evaluation on benchmark datasets of various scRNA-seq protocols, 10x Visium, and Slide-seq ST platforms. Moreover, we confirmed the integration capability of LANTSA to transfer cell annotation on large-scale and cross-platform scRNA-seq datasets. Finally, we validated the effectiveness of LANTSA for the identification of multiple mouse brain areas as well as the spatial mapping of cell types within cortical layers by integrating scRNA-seq and ST data.

## INTRODUCTION

Recent advances in transcriptomics technologies [1-4] have enabled high-throughput sequencing of mRNA at single-cell resolution or coupled with spatial information in multicellular organisms. Single-cell RNA sequencing (scRNA-seq) allows us to characterize the transcriptional landscape of individual cells in a way that provides novel insights [5, 6]. Spatial transcriptomics (ST) technologies resolve cellular localizations to unveil the organizational landscape of complex tissues [7, 8]. These cutting-edge technologies have begun to make important contributions to fundamental biological research and clinical applications [9, 10]. They have also promoted significant innovation in the computational methods devoted to deconvolve the biological implications underlying datasets.

Recently, numerous models have been proposed for the various tasks needed for scRNA-seq profiles, including clustering [11, 12], visualization [13], trajectory inference, and data integration [14]. Some of these tasks also provide manipulation techniques suitable for ST datasets [14, 15]. At the same time, several methods currently developed for ST profiles are applied to spatial clustering [16], or inference of cell types with the integration of scRNA-seq datasets [7, 17, 18]. For example, Seurat [14] and SCANPY [15], which accommodate multiple data processing approaches, provide comprehensive pipelines for both scRNA-seq and ST analyses. scVI [19] transforms data by a variational autoencoder (VAE) model, which provides favorable solutions for multiple tasks on scRNA-seq profiles. SC3 [20] aggregates Principal Component Analysis (PCA) and Laplace transformations, and then constructs a consensus matrix for cell clustering. Giotto [21], BayesSpace [16], and Hidden Markov Random Field (HMRF) [22] are all proposed for spatial clustering, which also incorporate spatial information to obtain a cluster assignment. In addition, SPOTlight [18], spatialDWLS [23], stereoscope [24] and robust cell-type decomposition (RCTD) [7] are specialized for spatial deconvolution by integrating spatial and single-cell transcriptomics datasets. The former two adopt a regression strategy (non-negative least squares regression for SPOTlight and dampened weighted least square regression for spatialDWLS) to deconvolve cell types in spatial locations and the latter two model a probabilistic distribution (negative binomial distribution for stereoscope and Poisson distribution for RCTD) to estimate the composition of spatial mixtures.

These methods have achieved many diverse applications but still have essential limitations: 1) Most of the methods assume that one (common) low dimensional space can mostly retain the biological variability underlying the original data, which is not justified due to the existence of multiple implicit subspaces underlying heterogeneous cell types or micro-environments [5, 19, 25]. 2) The analysis of large datasets or the performance of cross-platform integration requires a high level of computational capacities [14, 19] due to the expanding size of recent transcriptomics data. However, few of the existing methods can simultaneously fulfill the requirements. In particular, effective methods to integrate scRNA-seq and ST datasets for dissecting the spatial architecture into single cell types are still lacking [7].

We developed LANTSA, a landmark-based transferable subspace analysis framework, to overcome these limitations. LANTSA can separately or in an integrated way analyze scRNA-seq and ST data. The core model is adapted from subspace analysis theory, which has been proven to capture data structures in a more accurate way compared with traditional methods [26]. Specifically, LANTSA approximates the whole representation graph (i.e., sample-by-sample relationship) by representing each landmark sample as a linear combination of all samples based on a novel subspace model which preserves local structures. Thus, LANTSA can not only capture more accurate sample representation graph, but also significantly reduce the runtime and memory consumption. Subsequently, the representation graph could be interoperable with clustering methods (e.g., Leiden [28]) for identifying cell or spatial clusters, and dimensionality reduction methods (e.g., Uniform Manifold Approximation and Projection, UMAP [13]) for data visualization. In addition, LANTSA can also be applied for data integration via a learning and prediction process. LANTSA firstly relies on a linear dimensionality reduction approach based on a learning scRNA-seq dataset to obtain discriminants that best fit with the data representation configuration. Next, LANTSA builds a predictor with the learned discriminants and cell labels (derived from learning data), to annotate the queried scRNA-seq or ST profiles. In such a way, LANTSA can effectively analyze or integrate large datasets in an accurate and efficient manner.

We confirmed that LANTSA can accurately capture data structures as it outperformed state-of-the-art methods to identify cell and spatial clusters using a collection of scRNA-seq, 10x Visium, and Slide-seq V2 [29] datasets. We verified that LANTSA is approaching these fast dimensionality reduction-based methods in terms of time efficiency. We further demonstrated that the discriminant matrix of LANTSA showed great performance and suitability for label transfer across sequencing platforms on scRNA-seq and single-nucleus sequencing (snRNA-seq) cerebellum datasets. Finally, we conducted an in-depth study of a mouse brain Visium array, in which LANTSA successfully recognized brain architecture through spatial clustering, and deconvolved the cell mixtures by integrating an independent scRNA-seq dataset.

## MATERIAL AND METHODS

### Method Overview

LANTSA provides a computational framework for both separately and in an integrated way analyzing scRNA-seq and spatial transcriptomics (Figure 1 and Supplementary Figure S2 and S3). For individual datasets, LANTSA selects a small number of landmark samples ℒ and constructs a landmark-by-sample representation matrix *Z* by representing each landmark by other samples (Algorithm 1 in Supplementary Materials). There are two main advantages of such an implementation. The representation of samples to a limited number of landmarks instead of all samples 1) avoids the distraction by redundant or noisy samples (especially for massive-volume datasets); 2) significantly reduces both runtime (Figure 2B) and memory occupation due to the linear complexity of the solution (Section S1.2 in Supplementary Materials). In this regard, the representation matrix obtained by LANTSA accurately and efficiently characterizes the relationship of samples, which then serves for sample clustering and data visualization (Figure 1A). Note that the selection of landmarks is a crucial step to the downstream performances of LANTSA, which adopts the maximal subgradient violation strategy previously proposed by Matsushima, S. and Brbic, M. (27). The subsampled landmarks are theoretically proven to cover all the possible subspaces (e.g., cell groups) using this strategy, and each selected landmark appears to be a representative of its local ‘manifold’ (Supplementary Figure S4). Such that the representation approximation preserves exactly the data structures underlying heterogeneous subspaces. Then, the optimal landmark-by-sample representation matrix is constructed through a novel subspace model, which preserves global and local structures underlying zero-inflated scRNA-seq or spatial transcriptomics data. For multiple datasets, LANTSA adopts a transfer strategy to aggregate two or more datasets, in which LANTSA transfers cell labels from the reference to query datasets via a learning and prediction process (Figure 1B). In the learning stage, LANTSA, which is applied to the learning dataset of scRNA-seq, derives the low-dimensional discriminant matrix *P* from the representation matrix (Algorithm 2 in Supplementary Materials). In the prediction stage, the discriminant matrix allows LANTSA to project all samples into the same space, where label transfer is performed from the learning to prediction datasets. The key design in transfer strategy is to obtain the optimal projection in an iteratively alternating manner [30] that could best fit the quantitative representation of multiple cell groups. Therefore, the shared space retains biological signals underlying the cell differences, which enables label transfer within large-scale or cross-platform datasets. Note that the transfer is based on the optimal solution of the representation matrix, which suggests that the capability to characterize data structures is essential for LANTSA.

**Figure 1.**
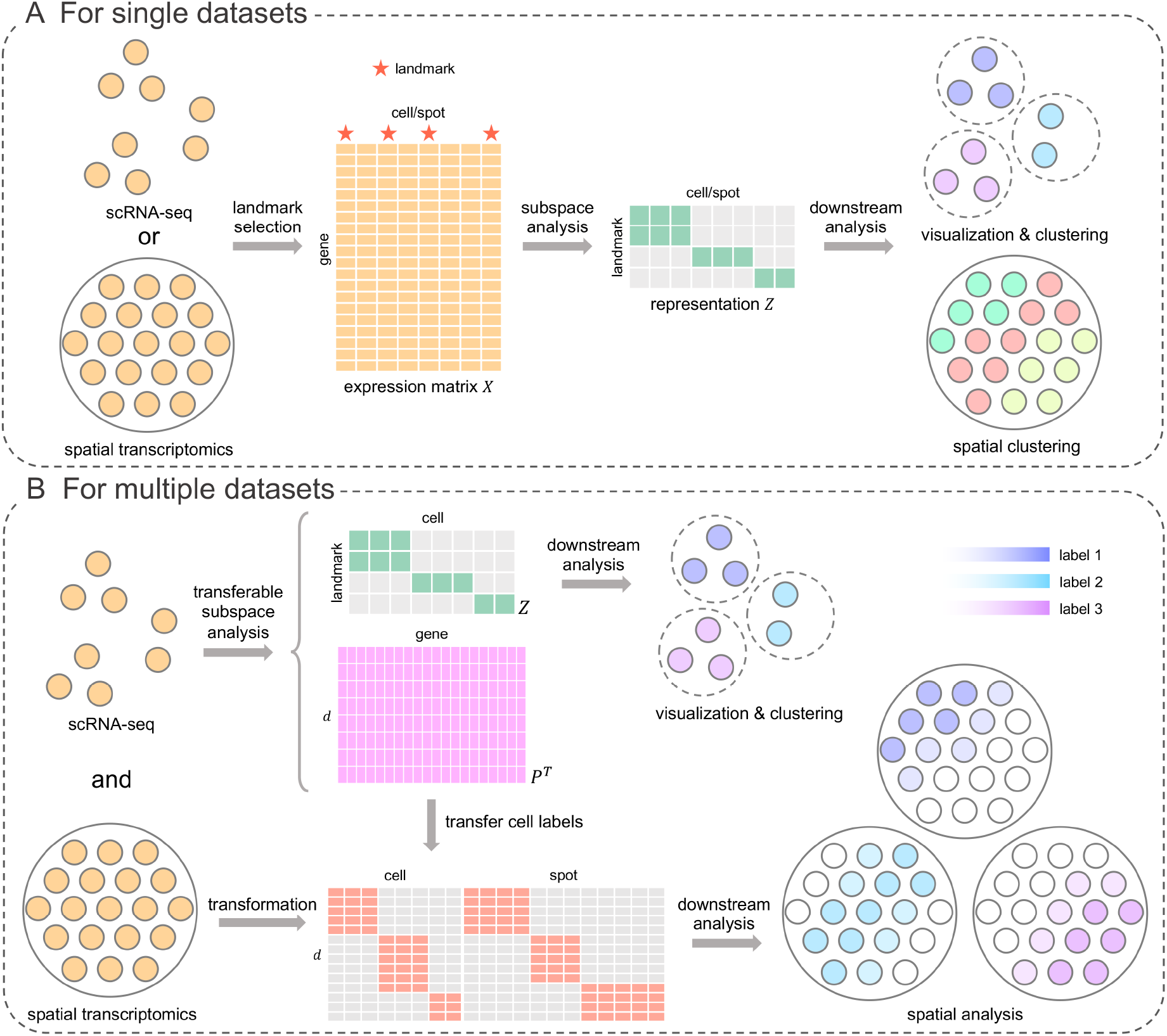
Illustrative scheme of LANTSA. (**A**) LANTSA for analyzing single (scRNA-seq or ST) datasets. Firstly, a set of landmark samples *ℒ* is selected from all samples (e.g., cells, spots) in the given transcriptomic dataset (see Methods). Next, the representation of landmarks to all samples (i.e., the matrix *Z*) is reconstructed through the sparse representation technique, which is regarded as the approximation of the relationship of all samples. Lastly, this representation matrix can be employed for various subsequent applications, e.g., UMAP visualization, identification of cell or spatial clusters. (**B**) Transfer strategy of LANTSA for integrating scRNA-seq and spatial transcriptomic profiles. The scRNA-seq (ST) data is denoted as the learning (prediction) dataset. From scRNA-seq data, LANTSA iteratively learns the optimal discriminant matrix *P* underlying the data structure (i.e., representation matrix *Z*) (see Methods). LANTSA transforms both learning and prediction datasets into the same space with the discriminant matrix, enabling the probabilistic transfer of cell labels from scRNA-seq data to the queried locations of ST data. Note that the model also allows label inference within large-scale or across scRNA-seq datasets. Each dot indicates the sequenced cell or spot. Dot color distinguishes cell/spatial groups after cluster analysis. Gradient color is specified for the inferred probability of a given cell type.

**Figure 2.**
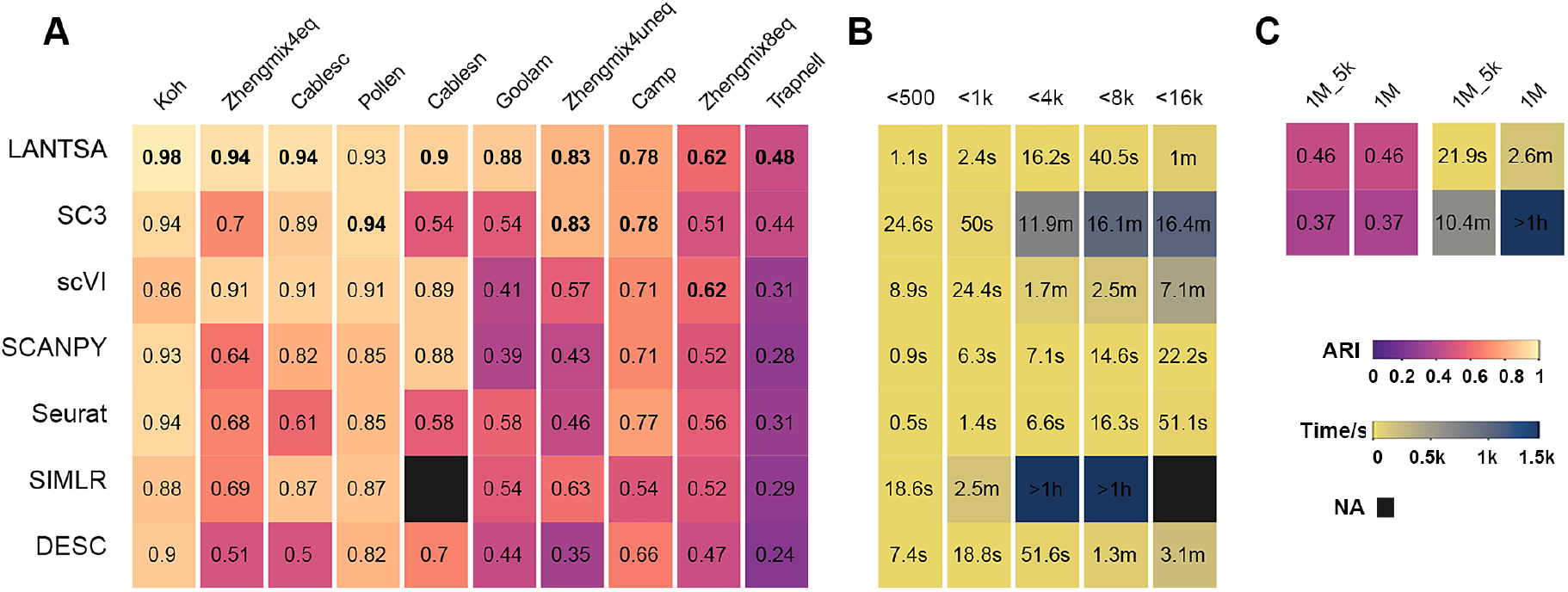
Clustering performance of LANTSA and the other six methods on scRNA-seq datasets. ARI values (**A**), running time (**B**) on 10 scRNA-seq datasets. Rows correspond to algorithms ordered by decreasing mean of per-dataset ARI values. Columns in (**A**) correspond to datasets ordered by decreasing ARI values of the performance of LANTSA. Columns in (**B**) correspond to categorized datasets based on sample size and each cell represents the average running time of this sample-size interval for every algorithm. Each cell displays the corresponding values and the highest ARI values for each dataset are highlighted in bold, except NAs. NA means that the algorithm cannot run on that dataset due to excessive memory footprint (using 128G RAM). The ARI values and running times are also calculated in repeated experiments on 1M dataset. These experiments involve the top two best methods, i.e., LANTSA and SC3; the average results are illustrated in (**C**). 1M_5k means the subset of 5,000 cells randomly selected from the 1M dataset, which is subsequently served as the learning set to predict the remaining cells (see Methods). The experiment was repeated 100 times and the detailed performance is shown in Supplementary Figure S10. Algorithms were tested on a computer with one 8-core Intel i9-9900K CPU addressing 128G RAM and one NVIDIA GeForce GTX 1660 GPU addressing 6G VRAM. Note that LANTSA, scVI and DESC support GPU acceleration. These methods were therefore assessed on GPU. Unit abbreviations: k, × 10^3^; s, second; m, minute; h, hour.

### LANTSA implementation for single datasets

#### Landmark sample selection by a violating subgradient strategy

A data matrix *X* = (*x*_1_, *x*_2_, … *x*_*N*_) ∈ *R*^*M*×*N*^ is given, where *M* and *N* denote the number of features (e.g., genes) and samples (e.g., cells, spots), respectively. Each sample can be represented by all samples according to traditional subspace analysis methods to capture the sample-by-sample relationship. The time and memory complexity of this solution is *O*(*N*^2^) (27), which is unacceptable for large-scale datasets. LANTSA proposes a critical assumption to overcome this limitation, namely that the sample-by-sample relationship is approximated only with a set of selected samples (i.e., landmark samples), instead of all samples. Thus, the formulation of LANTSA for representing landmark sample *i* is written as follows:

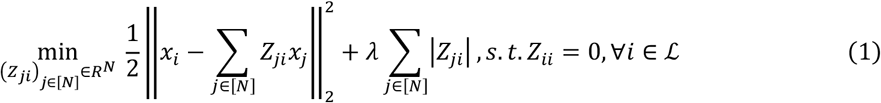

where [*N*] and ℒ represent the set of all samples and landmarks respectively, and *L* denotes the number of the landmark samples in ℒ with *L* << *N. Z*_,+_ denotes the representation strength between sample *j* and sample *i*, and a larger value indicates that two samples share more similarities. *λ* is a tunable parameter to balance the terms of representation error and sparsity of *Z*. Clearly, the time and memory complexity of this solution could be reduced to *O*(*LN*), which is significantly smaller than the original *O*(*N*^2^). The key problem now is the selection of samples to assemble ℒ which is assumed to cover all the subspaces underlying the data.

We leverage an incremental algorithm to obtain ℒ based on stochastic subgradient approximation to solve this problem according to Matsushima, S. and Brbic, M. [27], which theoretically guarantees that each landmark is a representative of its underlying subspace.

#### Capturing the relationship of samples based on landmark samples

After the landmark set ℒ is assembled, the representation matrix *Z* of samples can be solved by a novel subspace model (Algorithm 1 in Supplementary Material), which preserves the global and local structure underlying data. The formulation of LANTSA for capturing the representation matrix *Z* is written as follows:

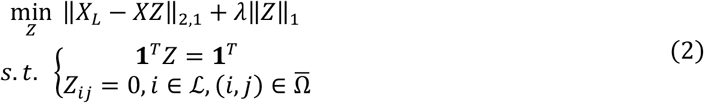

where *X*_*L*_ ∈ R^*M*×*L*^ is the expression matrix of landmark samples ℒ. **1** represents an all-one vector for normalization. 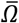 is the complement of *Ω* which is a set of connections of samples in a adjacency graph. For instance, if *x*_*i*;_ and *x*_*j*_ are not connected in the adjacency graph, then we have 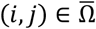. The adjacency graph is determined by *K*-nearest neighbor (*K*NN) algorithm with Euclidean distances of all samples on landmarks. Thus, the second constraint can guarantee the preservation of local structure underlying data. Parameter *K* may be chosen freely and there is a tunable parameter *λ* to balance the two terms in Eq. (2). Both parameters can be determined according to data properties or settled empirically.

The relationship of samples *Z* in the objective function Eq. (2) can be solved by the alternating direction method of multipliers (ADMM) [30] and its computational complexity is analyzed in Section S1 of the Supplementary Materials.

Note that despite we use the same landmark selection approach as Matsushima, S. and Brbic, M., we adopt a different algorithm solving the landmark-by-sample representation matrix rather than Lasso (27). Our solution can capture the global and local structures of data enabling superior performance on zero-inflated scRNA-seq data (Supplementary Figure S1).

### LANTSA implementation for multiple or large-scale datasets

We define the term ‘learning and prediction set/dataset’ for the circumstances that LANTSA is applied for multiple or large-scale datasets. For multiple datasets, LANTSA adopts a (often annotated scRNA-seq) dataset as the learning dataset for constructing both representation matrix and discriminant matrix. The remaining datasets serve as prediction datasets whose labels are transferred from the learning dataset based on the discriminant matrix. In contrast, for large-scale datasets, LANTSA divides the full dataset into a learning and prediction set. The former is utilized for deriving discriminant matrix and representation matrix that is used for partitioning clusters. And the latter is then labeled based on discriminant matrix and the cluster assignment of the former. Therefore, the relationship of samples and discriminant features are captured on the learning set, and the labels of the prediction set are obtained through classification.

#### 1) Learning stage

The model utilized to identify the sample representation and the discriminant matrix on the learning set is formulated as follows:

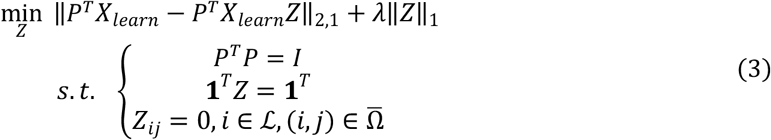

where *X*_*learn*_ is the expression matrix of learning samples. *Z*, which was previously defined as the representation matrix, reflects the relationship of samples. *P* denotes discriminant matrix which is a low-dimensional orthogonal matrix that contains the information of discriminative features affecting the relationship of samples. It can be used to construct a classification model for cell annotation on the prediction set.

Next, we focus on solving the objective function. An alternating iteration algorithm is designed to solve one matrix with the other fixed by considering that each term of the objective function can be solved separately. The computational procedure of the splitting problems is given in Section S1 of the Supplementary Materials. Note that this alternating solution guarantees an optimal discriminant matrix to smoothly fit with the overall structure of the representation matrix.

#### 2) Prediction stage

We use a set of discriminant vectors obtained from the learning set and the inferred labels from the representation matrix *Z* (or the prior known sample labels) to build a *K*NN classifier, which then transfers annotations to the prediction set. The pseudocode of the transfer strategy is summarized in Algorithm 2, which is stated in Section S1 of the Supplementary Materials.

### Clustering, visualization and classification approaches

LANTSA infers the cluster information by Leiden algorithm [28] given representation matrix *Z*. The parameter ‘resolution’ is adjusted to match the number of cell populations provided by the original authors or manually defined with prior (anatomical) knowledge.

LANTSA adopts the UMAP method for cell (spot) embedding visualization based on the representation matrix *Z*. In particular, the algorithm adopts a transfer strategy when processing large-scale data to reduce dimension based on discriminant matrix. It subsequently uses scanpy.pp.pca(), scanpy.pp.neighbors(), and then UMAP for visualization.

When transferring labels to prediction set (within or across platforms), we constructed a *K*NN classifier with the learned discriminant vectors and the clusters obtained from the learning dataset. Additionally, the prediction performance is impacted by the number of discriminant vectors. We picked the first *d* eigen vectors of the discriminant matrix *P* in the ascending order of eigen values. We used the KNeighborsClassifier class in the scikit-learn Python module for classification in our experiments with the parameter settings of ‘metric=cosine’ and ‘n_neighbors=10’. The predicted label for each sample was extracted by .predict() method, and the transferred probability for spot annotation was acquired from .predict_proba() method.

### Measurements of clustering and prediction performance

The clustering performance is quantified, if ground truth labels are available (e.g., from original publications), based on two widely used metrics: adjusted Rand index (ARI) [31] and normalized mutual information (NMI) [32]. We also used these two metrics for prediction accuracy by the label transfer approach allowing brief comparison of clustering and prediction outcomes.

### Identification of marker genes

Marker genes were identified for each spatial clusters in the 10x Visium sagittal array via ‘FindAllMarkers()’ in Seurat using default parameters.

We selected genes for each cell type in the reference cerebellum datasets with a minimum average expression above 1.5 × 10^−4^ counts and at least 1.25 log fold change (FC) compared to the average expression across all other cell types.

### Mapping bead clusters to reference cell types

Marker genes from reference cerebellum datasets help annotate bead clusters in Slide-seq data. We computed the FC score of each marker gene for given clusters over other beads. Next, we averaged the FC scores and mapped the cluster of the highest value to the corresponding single-cell cell type.

### Measurement of discriminant vectors

The capability of the discriminant matrix of LANTSA to preserve the global data structure and coherence of cell populations in the shared space is measured using different approaches.

We computed the cell-to-cell distance matrix in the original PCA and transferred spaces. Next, we computed the Pearson correlation of cell distances in two spaces for each cell. We binned the correlation values into 100 equal-width bins to facilitate visualization of this relationship and used a histogram plot to summarize the preservation of global structure in the data.

We used a metric of discrimination score (DS) first proposed in our previous work [5] to evaluate how the discriminants of LANTSA contribute to cluster separations. The metric is redefined as distant discrimination score (DDS) to measure the scaled distance between the most distant group pairs and neighboring discrimination score (NDS) for the neighboring groups. The range is from +1 for those cells that are tightly grouped in the feature space far from other clusters to −1 for very poor discriminative capability. In general, NDS is lower than DDS. If the NDS score is greater than 0, it indicates a positive contribution of the feature to distinguish the closest clusters.

### Functional enrichment analysis

The online platform of Metascape [33] was used for functional enrichment analysis of marker gene sets. We focused on the marker genes identified in three collections of cerebral cortices (layers 2—6), thalamus/hypothalamus and fiber tracts, which were sorted in ascending order of adjusted *P*-values in each collection and the top 300 marker genes were submitted for enrichment analysis. Enrichment analysis was carried out for each given gene list with the ontology sources from GO biological processes, KEGG pathway, Reactome gene sets and WikiPathways, using the default filtering thresholds. The results of the top 20 enriched terms were returned by Metascape as shown in Supplementary Figure S30, and five of these terms were selected for representative illustration in Figure 6B. The three analyzed segments were proven to have high proportions of excitatory neurons, inhibitory neurons and oligodendrocytes, according to the functional annotations.

## RESULTS

### Evaluation on single cell transcriptomics datasets

We firstly evaluated the ability of LANTSA to capture accurate relationship of cells on clustering of scRNA-seq data using ten real datasets spanning different technical and biological characteristics with annotations independent of benchmarked methods here (Table 1, Data availability, data preprocessing in Section S1.4). We benchmarked six popular methods originally designed for single-cell clustering or comprehensive analysis, i.e., SC3 [20], SCANPY [15], Seurat [14], Single-cell Interpretation via Multi-kernel Learning (SIMLR) [34], DESC [35] and scVI [19]. These methods were evaluated following their tutorials or using default parameters (Section S1.5 in Supplementary Materials). We used the adjusted Rand Index (ARI) and normalized mutual information (NMI) (see Methods) to quantify the similarity between reference labels and the clusters obtained by algorithms. The evaluation results illustrate that LANTSA generally performed better than other assessed methods on most of datasets (Wilcoxon signed-rank test, *P* < 0.01, Figure 2A and Supplementary Figure S5). Data visualization in UMAP embeddings qualitatively supported the clustering performance (Supplementary Figure S6 and S7). In addition to accuracy, we also compared the running time on these datasets. The dimension-reduction-first methods, i.e., SCANPY and Seurat, were demonstrated to be the most time-efficient (Supplementary Figure S8). LANTSA also impressively took less than one minute when processing less than 16,000 cells (Figure 2B) even though it does not employ dimension reduction.

**Table 1.** Single cell transcriptomic datasets for LANTSA evaluation. Note that the ground truth labels of all datasets are independent of the benchmarked methods here, which are listed at Data availability. k, number of groups in ground truth labels; *N*, total number of cells.

| Dataset | $k$ | $N$ | Unit | Description | Protocol | Reference |
| --- | --- | --- | --- | --- | --- | --- |
| Camp | 7 | 777 | FPKM | Human liver bud | SMARTer | Ref. [50] |
| Goolam | 5 | 124 | CPM | Mouse embryo | Smart-Seq2 | Ref. [51] |
| Pollen | 11 | 301 | TPM | Human cell line | SMARTer | Ref. [52] |
| Koh | 9 | 531 | Count | FACS purified H7 human embryonic stem cells | SMARTer | Ref. [53] |
| Trapnell | 3 | 222 | Count | Human skeletal muscle myoblast cells | SMARTer | Ref. [54] |
| Zhengmix4eq | 4 | 3,994 | Count | Mixture of purified peripheral blood mononuclear cells | 10x Genomics GemCode | Ref. [55] |
| Zhengmix4uneq | 4 | 6,598 | Count | Mixture of purified peripheral blood mononuclear cells | 10x Genomics GemCode | Ref. [55] |
| Zhengmix8eq | 8 | 3,994 | Count | Mixture of purified peripheral blood mononuclear cells | 10x Genomics GemCode | Ref. [55] |
| Cablesc | 11 | 4,857 | Count | Mouse cerebellum | Drop-seq | Ref. [7] |
| Cablesn | 19 | 15,609 | Count | Mouse cerebellum | 10x single-nucleus RNA-seq | Ref. [7] |
| 1M | 20 | 1,306,127 | Count | Mouse brain | 10x Genomics GemCode | Ref. [56] |

**Table 2.** Number of marker genes for specific cell types obtained in cerebellum datasets

| **Dataset** | **Astrocytes** | **Bergmann** | **Choroid** | **Granule** | **Oligodendrocytes** | **Purkinje** | **MLI** |
| --- | --- | --- | --- | --- | --- | --- | --- |
| **Cablesc** | **144** | **175** | **226** | **144** | **236** | **309** | **-** |
| **Cablesn** | **180** | **135** | **181** | **147** | **284** | **167** | **104** |

Next, we used another scRNA-seq dataset of 1.3 million mouse brain cells (i.e., 1M) (Table 1), to assess the capability to process large datasets. We adopted the transfer strategy of LANTSA to consider the possible consumption of time and memory and benchmarked against SC3 that also utilizes a similar hybrid strategy for large-scale datasets [20]. We randomly selected 5,000 cells as the learning set. LANTSA identified the cell groups of this subset and obtained discriminant features, which were subsequently used to train a *K*NN classifier to predict labels of the remaining cells (see Methods). Repeated results for 100 tests show high correlation between clustering (for the learning set) and classification performance (for the prediction set) (Supplementary Figure S9), which verifies the feasibility of our transfer strategy. LANTSA outperformed SC3 using a similar process in terms of clustering accuracy and running time (Figure 2C and Supplementary Figure S10). In fact, LANTSA obtains the discriminant features depending only on a few optimization iterations (see Methods), and it could considerably save on the prediction time.

### Performance of LANTSA across various parameter settings

There are mainly three tunable parameters in LANTSA implementation, i.e., *λ*, the number of neighbors and the number of landmarks. *λ* is a regularization parameter, which controls the sparsity of sample representation. The number of neighbors denotes how many neighbors to use for constructing a *K*NN graph that serves as a hard constraint in subspace analysis, which is supposed to obtain a more coherent representation. The number of landmarks determines how many samples to be selected as representatives of the implicit subspaces or sample clusters, which balances the accuracy and efficiency.

We performed sensitivity analysis of these three critical parameters on Koh (531 cells) and Zhengmix4eq (3,994 cells) datasets. We showed that the clustering performance of LANTSA is robust to a wide range of parameter settings (Supplementary Figure S11). Furthermore, we ran LANTSA with 100 different random seeds to test its stability under various number of selected landmarks. The results indicated that the performance of LANTSA will reach a “plateau” when adequate landmarks were selected to cover possible subspaces or sample clusters, and LANTSA is insensitive to the selected landmark size (standard deviation varies from 0.010 to 0.045) and identified accurate clusters (Supplementary Figure S11C) after reaching the “plateau”. However, excessive landmarks may increase time and memory overhead especially for large-scale datasets. We evaluated the time efficiency under various number of landmarks on two relatively large datasets (Zhengmix4eq with 3,994 cells and Zhengmix4uneq with 6,498 cells), and the results illustrate that time consumption is linear to the number of landmarks for both datasets (Supplementary Figure S12, adjusted R-squared 0.97 for Zhengmix4eq and 0.99 for Zhengmix4uneq, *F* test *P* < 10^−15^ for both datasets). We generally suggest that number of landmarks should be set to 20-30 times the empirical number of clusters, which still occupy a small proportion of entire samples, to achieve remarkable accuracy and computational efficiency.

### Evaluation on 10x Visium spatial transcriptomics datasets

We next evaluated the ability of LANTSA to identify spot clusters using 14 10x Visium arrays: 12 dorsolateral prefrontal cortex and 2 mouse brain sagittal sections (data preprocessing in Section S1.4). Here, we considered six methods for comparison, including SC3, SCANPY, Seurat and three methods specialized for processing spatial transcriptomics, i.e., BayesSpace [16], Giotto [21] and stLearn [17]. These methods were evaluated following their tutorials or using default parameters (Section S1.6 in Supplementary Materials). We used ARI to quantify the similarity between identified clusters and manual annotations which are provided by spatialLIBD package [36] for 12 cortex samples. For other two sagittal samples, we assessed the clusters *in situ* with the neuroanatomical annotated images obtained from the Allen Mouse Brain Atlas [37] as the ground truth reference.

The quantitative results show that spatial clustering methods generally perform better than non-spatial methods (Figure 3A, Wilcoxon signed-rank test, *P* < 10^−4^), since they incorporate spatial information into cluster inference. LANTSA was demonstrated to mostlyoutperform other partitions from non-spatial or spatial clustering methods (Figure 3A, Wilcoxon signed-rank test, *P* < 10^−6^). BayesSpace could also provide clear spatial separations, while its accurate probabilistic inference consumes more time. In fact, LANTSA has obvious advantage in time efficiency over most of the involved methods except SCANPY (Figure 3B). In addition, the qualitative comparison shows that LANTSA could excel at distinguishing subtle structures in mouse brain arrays (Figure 3, UMAP visualization in Supplementary Figure S13). According to reference maps, LANTSA identified the cerebral cortex (CTX) and lateral ventricle (VL) section in the anterior slice (Figure 3C), the dentate gyrus (DG), and cornu ammonis (CA) sections of the hippocampus region in the posterior slice (Figure 3D). Other benchmarked methods failed to localize the fine structures or only identified fewer sections.

**Figure 3.**
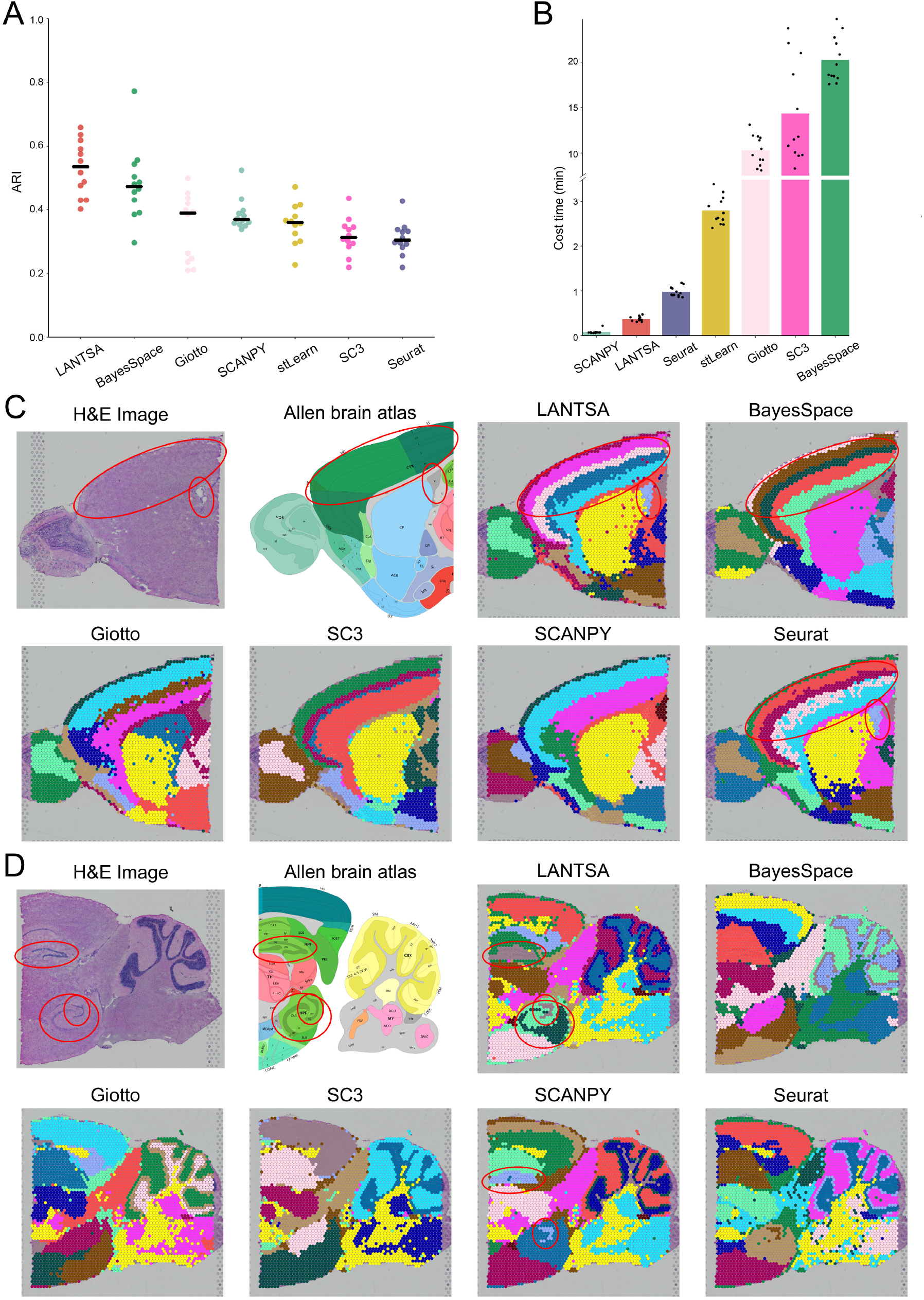
Comparative performance of LANTSA to existing spatial and non-spatial methods for spatial clustering. Summary of clustering performance on 12 manually annotated spatialLIBD datasets in terms of ARI values (**A**) and running time (**B**). The ground truth labels for these datasets can be found in the original publication. Each point denotes the measured performance on one dataset. The center line in (**A**) indicates the median ARI value of each method on all datasets. The methods in (**A**) are ordered by decreasing median ARI values. The height of bar in (**B**) indicates the mean running time of each method on all datasets. The methods in (**B**) are ordered by increasing mean running time. The spatial clustering of mouse brain sagittal anterior (**C**) and posterior (**D**) slices. The hematoxylin and eosin (H&E) images of each dataset (first images in **C** and **D**) and the corresponding anatomical definitions obtained from the Allen Mouse Brain Atlas (second images in **C** and **D**) are shown as references. The identified clusters by all the involved approaches are illustrated on H&E staining images and distinguished using different colors without anatomical correspondence. Fine anatomical regions, for example CTX, VL sections in (**C**) and DG, CA sections in (**D**) are marked by red circles on reference images and computational results (if any exists).

### LANTSA enables transfer of annotations across platforms

The two-step strategy (i.e., first clustering then prediction) of LANTSA is primarily tailored for large datasets (e.g., 1M). Here, we benchmarked the cross-dataset prediction performance of LANTSA on snRNA-seq and scRNA-seq cerebellum datasets through bidirectional predictions (see Methods), along with six methods (i.e., RCTD [7], scanorama [40], Seurat [41], spatialDWLS [23], SPOTlight [18] and stereoscope [24]) capable of label transfer across platforms (implementation details in Section S1.7). The predication outcomes of LANTSA are more accurate with the prediction accuracy of 0.97 on scRNA-seq dataset and 0.95 on snRNA-seq (Supplementary Figure S23), which implies the insensitivity of LANTSA to reference platforms. Cross-prediction could well distinguish endothelial cells, granule cells, choroid cells, Purkinje cells, oligodendrocytes and Bergmann cells for both datasets (Figure 4A and 4B) (prediction proportion ≥ 0.9). For scRNA-seq data, the choroid group is shown to be divided into two subgroups no matter in raw data space (Supplementary Figure S24A) or discriminant space (Figure 4C), which probably results in more mispredictions. A small number of oligodendrocytes, microglia are mixed with granule cells, which also decreases the transfer performance. Bergmann cell, a specialized astroglia [42], was biologically similar to astrocytes for snRNA-seq data (Figure 4D and Supplementary Figure S24B). Thus, LANTSA did not correctly label a few cells in these two groups. The overall observations from UMAP visualizations (Figure 4C, 4D and Supplementary Figure S25 and S26) suggest that our transfer strategy more likely retains biological structures from the learned dataset, rather than correction of batch effects.

**Figure 4.**
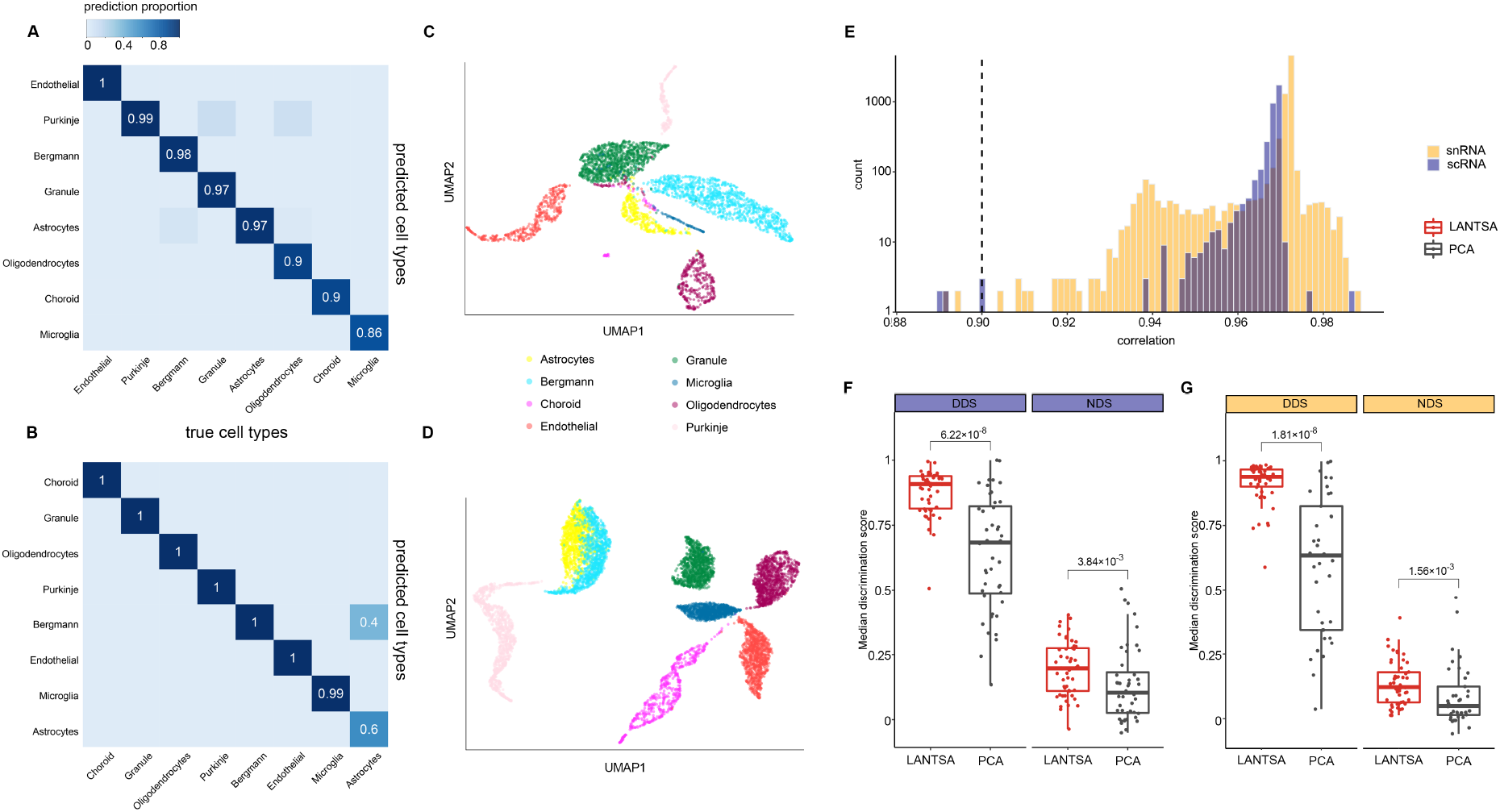
LANTSA transfers biological information between snRNA-seq and scRNA-seq cerebellum datasets. Confusion matrix of true versus predicted cell types across platforms either learned from snRNA-seq and predicted on scRNA-seq data (**A**) or in reverse (**B**). Color represents the proportion of the cell type on the *x* axis predicted as the cell type on the *y* axis. UMAP visualization of cells projected from LANTSA discriminant space in prediction set: scRNA-seq data (**C**) and snRNA-seq data (**D**). Color represents the prior cell annotations provided from the original paper. (**E**) Distributions of Pearson correlation values between cell distances in LANTSA-transferred space and original PCA space. Log10 transformed *y* axis indicates the number of cells that fall in each correlation interval (i.e., *x* axis). Dashed line marks correlation of 0.90. Discriminative abilities of the first 50 discriminant vectors (vs. first 50 principal components for PCA) to separate cell populations between the most distant pairs (**F** and **G** left) or neighboring groups (**F** and **G** right). The navy bars in E and F represent evaluation for the scRNA-seq cerebellum dataset; and the orange bars in E and G for snRNA-seq dataset. DDS, discrimination score for distant groups; NDS, discrimination score for neighboring groups (as defined in [5]). Discrimination scores smaller than zero are not shown. When the score is closer to 1, the discrimination ability is stronger for that vector or component. Wilcoxon signed-rank test is used to test the difference between discrimination score distributions. The obtained *P*-values are corrected by FDR and labeled on the box top.

We further investigated the shared space constructed by LANTSA. First, we analyzed whether the ‘transferred’ projection preserves global data structure as in the original dataset. We compared the distance preservation in low-dimensional embeddings for each cell (see Methods) (Supplementary Figure S27 and S28) and found a large proportion of cells in the transferred space that consistently kept the distances in the original (i.e., PCA) embeddings (the distance correlation reached 0.90 for more than 99.5% cells) (Figure 4E). Furthermore, we used the discrimination score, a score between −1 (for the worst) and 1 (for the best), to investigate the ability of loadings to separate different cell types (see Methods). The significant score differences revealed that the discriminant vectors of LANTSA appeared more capable of distinguishing cell populations than the principal components from PCA (Figure 4F and 4G, Wilcoxon signed-rank test, *P* < 0.01). Altogether, these results illustrate two key characteristics of our transfer matrix (i.e., preserving global data structure and separating different cell types), which contributes to its excellent ability to deal with large-scale or cross-platform datasets.

### LANTSA localizes fine spatial features in 10x cerebrum data

Finally, we moved on to an in-depth 10x Visium case for validating the capability of LANTSA to cope with spatial transcriptomics datasets. We named the clusters of sagittal anterior slice using the Allen taxonomy as reference (Supplementary Figure S29), which was previously identified in the benchmark study (Figure 3C). We searched for the differentially expressed marker genes of all the sections (see Methods). The top 1–3 marker genes showed strong specificity between different spatial structures (Figure 5A). The cortical layer 1 highly expresses *Ptgds*, a highly enriched gene in the mouse meninges [38], which is different from layers 2–6 that specifically express neuron-related genes, e.g., *Nrgn, Cck*, and *Lamp5* [43]. Fiber tracts express a high level of the myelin proteins *Plp1, Mobp* and *Mbp*, which indicates they may accommodate many oligodendrocytes [38]. The inhibitory neuron markers *Pvalb* and *Sst* [44] are abundant in the thalamus and hypothalamus. We next broadly classified functional modules of the top 300 differentially expressed genes in the regions of the cerebral cortex (layer 1 excluded), thalamus/hypothalamus, and fiber tracts. These top enriched functional annotations could indeed roughly separate these structures into distinct categories of abundant excitatory neurons (associated with glutamatergic neurotransmitter), inhibitory neurons (dominated with GABAergic synaptic transmission), and oligodendrocytes (involved in myelin related processes) (Figure 5B and Supplementary Figure S30) [45].

**Figure 5.**
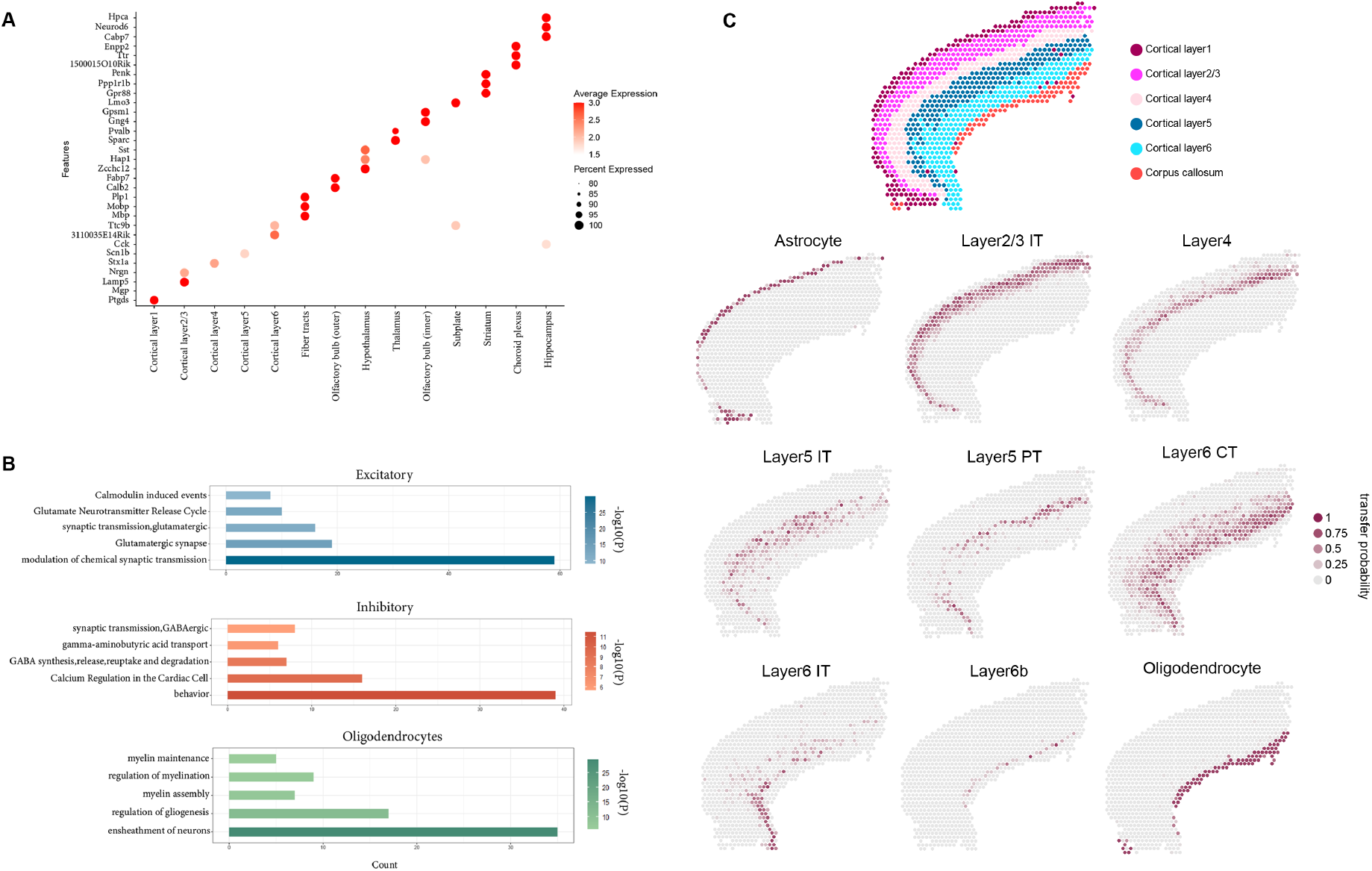
LANTSA localizes fine spatial features in brain sagittal anterior section. (**A**) Bubble chart of marker genes of identified anatomical regions. The size of each circle reflects the percentage of cells in a cluster where the gene expression is detected, and the color intensity reflects the average expression level within each cluster. (**B**) Curated functional terms enriched in three collections of differentially expressed genes. The excitatory category sweeps marker genes of neuron layers (layers 2–6) in cerebral cortex, inhibitory category contains the union sets of thalamus and hypothalamus, and oligodendrocytes refers to those fiber tracts. The *x* axis represents the number of overlapped genes between query gene set and given ontology term. ‘log10(*P*)’ denotes the *P*-value in log base 10 and color density indicates enrichment significance. (**C**) Probabilistic transfer from scRNA-seq to sub-assay of cortex layers 1–6 and corpus callosum (part of fiber tracts). The first image in (**C**) is colored as shown in Supplementary Figure S29 and others are colored by transfer probability of specific cell type assigned to each spot. The three neuron types include: IT: intratelencephalic; PT: pyramidal tract; CT: corticothalamic cells.

Nevertheless, the spot diameter of Visium arrays is around 50-100 μm, which suggests the spot could be a mixture of multiple cell types [7]. Thus, we leveraged the transfer strategy to deconvolute the spatial contents from a mouse cortical scRNA-seq dataset. We performed probabilistic transfer for each scRNA-seq derived cell type (see Methods) in addition to the evaluation of cross-platform prediction. On the query subarray, LANTSA provides a more spatially coherent spot deconvolution, which exhibits distinct layers of cell types, compared to other involved methods (Supplementary Figure S31). For example, oligodendrocytes are exactly inferred to be mainly localized in the corpus callosum (part of the fiber tracts) (Figure 5C), which is consistent with the previous annotation of spatial structures (Figure 5A). Non-neuron layer 1 is predicted to be enriched for astrocytes as is generally accepted [46]. In addition, fine neuron types have a well-organized distribution across different cortex layers in agreement with our previously identified sections (Figure 5C). Intratelencephalic (IT) cells constitute the largest proportion of the three types of neurons and correspond well to all layers (layers 2–6) [47], and layer-specific IT types are appropriately separated (Figure 5C). While corticothalamic (CT) neurons are mainly distributed in layer 6a [48], at the bottom of which the thin layer 6b, is also identified by LANTSA. The results revealed that the discriminant features learnt from the cell reference graph can accurately confirm the regional enrichment of specific cell types on the expected structures of brain anatomy.

## DISCUSSION

Single-cell and spatial RNA measurements provide new perspectives to define biological heterogeneities of tissues and diseases. These approaches demand computational methods applied to a range of rather than individual analysis tasks [19]. However, the fundamental issue is to accurately capture the relationship of samples (e.g., cells, locations). At the same time, the current methods still need to be improved, especially for analyzing large-volume or cross-platform datasets. In this work, LANTSA, was designed as such a scalable framework that accounts for these requirements. LANTSA provides a computationally efficient tool to obtain representations of samples (i.e., sample-sample relationships) and features (i.e., transferable discriminant vectors), which are then used for accurate downstream analysis, including clustering, visualization, and data integration.

Specifically, LANTSA solves a representation graph by the method of selective sampling-based sparse subspace analysis [27], which accurately and efficiently captures the underlying data structures. In addition, LANTSA can serve as an integration method: it initially learns an optimal projection with the objective of maximizing the preservation of the representation structure, and then maps the query samples by combining the reference dataset into the shared space for label transfer. We benchmarked LANTSA with frequently used methods for clustering and data visualization on scRNA-seq and ST datasets to evaluate the representation of samples. We showed that LANTSA is comparable to the state-of-the-art methods for the identification of sample groups with a better balance of accuracy and efficiency. We validated the properties of LANTSA for the transfer matrix that enables data projection onto a low-dimensional space. The properties examined include the preservation of global data structure and the ability to separate different cell groups, which contribute to the transfer of annotation from a reference to unlabeled datasets. We successfully applied LANTSA to a large-volume dataset (e.g., 1M), single-cell RNA profiles of different platforms, and the integration of scRNA-seq and ST datasets using the transfer strategy. Although the data structure captured by LANTSA outperforms state-of-the-art methods in terms of accuracy and computational efficiency, LANTSA cannot track which functional gene module contributes to each sample group. Like current analysis methods, LANTSA mainly focused on the representation of samples rather than bridging the correspondence of sample and feature representations, which is a limitation of LANTSA for application to the in-depth analysis of biological interpretations. Certainly, most methods are compromised to use a separate manner: first to confirm the sample structure and subsequently to perform statistical tests for marker identification (or vice versa). However, the splitting solution may cause untraceable biological variabilities, even leading to biased representation of outcomes. Therefore, further study of the joint analysis of sample and feature representations is warranted.

## DATA AVAILABILITY

Python source code of LANTSA, under the open-source BSD 3-Clause license, is available at https://github.com/zccqq/LANTSA. The documentation website provides the installation guide, tutorials, and API references, which are available at https://lantsa.readthedocs.io/. LANTSA is also published as a Python package named ‘lantsa’ on Python Package Index (PyPI) at https://pypi.org/project/lantsa/ and can be directly installed via the pip installer.

The datasets used in single-cell clustering benchmark are available from original publications, which are listed in Table 1. The ground truth labels of Camp, Goolam and Pollen were previously used in a method publication [20], which are available at https://hemberg-lab.github.io/scRNA.seq.datasets/. The ground truth labels of Koh, Trapnell, Zhengmix4eq, Zhengmix4uneq and Zhengmix8eq can be found in a benchmark publication [49]. The ground truth labels of Cablesc and Cablesn were previously used in a method publication [7], which are available at https://singlecell.broadinstitute.org/single_cell/study/SCP948. The ground truth label of 1M is derived from Cell Ranger pipeline, which is available at https://singlecell.broadinstitute.org/single_cell/study/SCP383.

## ACKNOWLEDGEMENT

We thank the anonymous reviewers for useful suggestions. Q.S. and C.Z conceived and designed the framework and the experiments. L.W. and L.X. performed the experiments. L.W. developed the Python package and documentation website of the framework. L.W., Q.S. and C.Z analyzed the data and wrote the paper. W.G. and L.C. revised the manuscript.

## KEY POINTS

- We introduce subspace analysis approach to single-cell and spatial transcriptomics analysis. Traditional methods mostly follow a dimension-reduction-first schema, i.e., assume to map samples into one common lower-dimensional space, where the accurate data structures can be hardly captured due to the existence of possible subspaces underlying the data. LANTSA directly performs subspace analysis on the original dataset rather than the dimension-reduced latent space, which enables better characterization of sample-by-sample relationship.
- We propose landmark-based subspace analysis to deal with large-scale datasets. Expanding size in recent transcriptomics data proposes high demand for computation resources. LANTSA learns the representation of samples to landmarks (consisting of a small number of samples) to approximate the entire sample-by-sample representation, which saves time consumption and memory occupation to a great extent. Furthermore, the approach to select landmarks, i.e., maximal subgradient violation strategy, theoretically guarantees that these samples could cover all possible subspaces underlying data, which is also experimentally validated.
- We extend LANTSA with integration (i.e., transfer) strategy for analyzing large-scale and cross-platform datasets. Specifically, LANTSA captures discriminant matrix from one learning (often annotated scRNA-seq) dataset, and then projects the samples of all datasets into the discriminant space, where the labels (e.g., cell types) from learning dataset can be transferred to the unannotated samples. In particular, integration of spatial and single-cell transcriptomics datasets combines the merits from both technologies to unveil heterogeneous landscape of cellular organization within tissues.

## FUNDING

This work is supported by National Natural Science Foundation of China (Grant Nos. 61802141, 31930022, 31771476 and 62002329), Strategic Priority Research Program of the Chinese Academy of Sciences (Grant No. XDB38040400), Key scientific and technological projects of Henan Province (212102310083) and Henan postdoctoral foundation (202002021).

**Chuanchao Zhang** received the PHD from Wuhan University, Wuhan, China, in 2017. He is currently an assistant research fellow in Key Laboratory of Systems Health Science of Zhejiang Province, Hangzhou Institute for Advanced Study, University of Chinese Academy of Sciences, Chinese Academy of Sciences, Hangzhou 310024, China. His current research interests include bioinformatics, machine learning, single-cell transcriptomics and spatial transcriptomics.

**Lequn Wang** is a PhD student in the Center for Excellence in Molecular Cell Science, State Key Laboratory of Cell Biology, Shanghai Institute of Biochemistry and Cell Biology, Chinese Academy of Sciences, Shanghai 200031, China. His research interests include bioinformatics, machine learning, single-cell transcriptomics and spatial transcriptomics.

**Xinxing Li** is a master student in College of Informatics, Huazhong Agricultural University, Wuhan, China. Her current research interests include bioinformatics, machine learning, single-cell transcriptomics and spatial transcriptomics.

**Wei-Feng Guo** is an associate professor at the department of Electrical Engineering, Zheng Zhou university. Guo obtained the PhD degree in 2020 at the department of automation from Northwestern polytechtical university, China. Guo does researches in design of complex network control algorithms and its applications to human cancer genomics.

**Qianqian Shi** received the PHD from Shanghai Institute of Biological Sciences, University of Chinese Academy of Sciences, Chinese Academy of Sciences, China, in 2017. She is currently an associate professor at College of Informatics, Huazhong Agricultural University, Wuhan, China. Her current research interests include bioinformatics, machine learning, single-cell transcriptomics and spatial transcriptomics.

**Luonan Chen** is a professor and executive director in the Key Laboratory of Systems Biology, Shanghai Institute of Biochemistry and Cell Biology, Center for Excellence in Molecular Cell Science, Chinese Academy of Sciences, Shanghai 200031, China; Hangzhou Institute for Advanced Study, University of Chinese Academy of Sciences, Chinese Academy of Sciences, Hangzhou 310024, China; and Center for Excellence in Animal Evolution and Genetics, Chinese Academy of Sciences, Kunming 650223 China. His interests include systems biology, computational biology, bioinformatics and applied mathematics.

## Notes

### Competing Interest Statement

The authors have declared no competing interest.

